# Targeted re-sequencing reveals the genomic signatures of multiple barley domestications

**DOI:** 10.1101/070078

**Authors:** Artem Pankin, Janine Altmüller, Christian Becker, Maria von Korff

## Abstract

- Barley (*Hordeum vulgare* L.) is an established model to study domestication of the Fertile Crescent cereals. Recent molecular data suggested that domesticated barley genomes consist of the ancestral blocks descending from multiple wild barley populations. However, the relationship between the mosaic ancestry patterns and the process of domestication itself remained unclear.
- To address this knowledge gap, we identified candidate domestication genes using selection scans based on targeted resequencing of 433 wild and domesticated barley accessions. We conducted phylogenetic, population structure, and ancestry analyses to investigate the origin of the domesticated barley haplotypes separately at the neutral and candidate domestication loci.
- We discovered multiple selective sweeps that occurred on all barley chromosomes during domestication in the background of several ancestral wild populations. The ancestry analyses demonstrated that, although the ancestral blocks of the domesticated barley genomes descended from all over the Fertile Crescent, the candidate domestication loci originated specifically in its eastern and western parts.
- These findings provided first molecular evidence in favor of multiple barley domestications in the Levantine and Zagros clusters of the origin of agriculture.

## Introduction

Domesticated barley (*Hordeum vulgare* ssp. *vulgare*) is one of the Neolithic founder crops, which facilitated establishment of the early agricultural societies (Lev-Yadun *et al.*, 2000). Due to its striking environmental plasticity, barley is an important staple crop in a wide range of agricultural environments (Dawson *et al.*, 2015). The first traces of barley cultivation were found at archaeological sites in the Fertile Crescent, which dated back to ∼10,000 B.C. (Zohary *et al.*, 2012). The Fertile Crescent is the primary habitat of the crop progenitor wild barley (*H. vulgare* ssp. *spontaneum*). However, its isolated populations have spread as far as North African and European shores of the Mediterranean Basin and East Asia (Harlan and Zohary 1966). Wild barley is a rich yet underutilized reservoir of novel alleles for breeding of barley cultivars better adapted to predicted future climatic perturbations.

In contrast to some other crops, the visible phenotype of domesticated barley did not dramatically diverge from its wild form. So far, the spike rachis brittleness has remained the only well-characterized domestication trait that exhibits a clear dimorphism between the wild and domesticated subgroups, which are characterized by the brittle and non-brittle spikes, respectively (Abbo *et al.*, 2014; Pourkheirandish *et al.*, 2015). Other traits differentiated between the modern-day wild and domesticated genotypes and underlying genes that define the barley domestication syndrome, as a complex of all characters that characterize the domesticated phenotype, are yet undiscovered (Hammer, 1984; Meyer and Purugganan, 2013). When adaptive phenotypes are not clearly defined, the so-called bottom-up approach, which starts with the identification of genome-wide signatures of selection, has proven instrumental in reconstructing the genetic architecture of the domestication syndrome (Ross-Ibarra *et al.*, 2007; Shi and Lai 2015). In other crops, the selection scans detected multiple selective sweep regions associated with domestication, which comprised hundreds of candidate domestication genes (Huang *et al.*, 2012; Hufford *et al.*, 2012; Lin *et al.*, 2014; Schmutz *et al.*, 2014; Zhou *et al.*, 2015).

The circumstances of barley domestication are debatable and its genome-wide effects on the domesticated barley genomes remain poorly understood (Pankin and Korff, 2017). The early models based on diversity analyses of isolated genes and neutral DNA markers proposed the Israel-Jordan area as a primary center of cultivated barley origin and proposed the eastern Fertile Crescent, the Horn of Africa, Morocco and Tibet as the alternative centers of domestication (Negassa, 1985; Molina-Cano *et al.*, 1999; Badr *et al.*, 2000; Morrell and Clegg, 2007; Dai *et al.*, 2012). As regards the number and the timescale of domestication events, one school of thought maintains that Neolithic domestication in the Near East has been a rapid centric innovation (Abbo *et al.*, 2010; Heun *et al.*, 2012; Zohary *et al.*, 2012). Conversely, the archaeobotanical evidence and simulation studies prompted development of the so-called protracted domestication model, which postulates that domestication of the Near Eastern crops has been a slow polyphyletic process dispersed over large territories (Allaby *et al.*, 2008; Fuller *et al.*, 2011, 2012; Purugganan and Fuller 2011). The diphyletic origin of the non-brittle spike phenotype of cultivated barley and the heterogeneous (mosaic) ancestry of cultivated barley genomes, which consisted of ancestral fragments originating from several wild barley populations, supported the protracted model (Allaby, 2015; Poets *et al.*, 2015; Pourkheirandish *et al.*, 2015). The heterogeneous origin of domesticated barley genomes hints at the existence of several independent lineages of barley cultivation. However, the link between the mosaic ancestry patterns and the process of domestication remained unclear.

Here, we partitioned the domesticated barley genomes into the neutral (in relation to domestication) and domestication sweep regions identified by selection scans and separately reconstructed the phylogeographic history of their origin. To this end, we resequenced a diversity panel comprising 344 wild and 89 domesticated barley genotypes using a custom genome-wide enrichment assay (∼ 544,000 SNPs). The selections scans identified multiple domestication sweep regions on every barley chromosome. Analysis of the top candidate genes within the domestication sweeps suggested cases of parallelism in targets of selection during domestication of barley and other crop species. The patterns of ancestry at the neutral loci revealed signatures of abundant continuous gene flow, which hindered identification of lineages descending from the independent founder events. Nevertheless, heterogeneous ancestry of the domestication sweep loci provided the first molecular evidence that multiple domestication sweeps occurred in the background of several wild barley populations residing in the eastern and western clusters of the Fertile Crescent.

## Materials and methods

### Plant material and Btr genotyping assay

A panel consisting of 344 wild and 89 domesticated lines were selected to maximize genetic diversity and to cover the entire range of the wild and landrace barley habitats in the Fertile Crescent (**Supplementary table 1**). The elite barley cultivars were sampled to represent Northern European, East Asian, North American and Australian breeding programs. The largest part of the germplasm set, 98% of wild and 40% of domesticated barley genotypes originated from the area of the Fertile Crescent. The selection of domesticated barley originated from various breeding programs and represented the whole variety of cultivated barley lifeforms, namely two-(71%) and six-row (29%) genotypes with winter (45%) and spring (55%) growth habits based on the passport data. All material was purified by single-seed descent to eliminate accession heterogeneity.

Leaf samples for DNA extraction were collected from single 3-week old plants. The DNA was extracted using the DNeasy Plant Mini kit (QIAGEN, Hilden, Germany) and quantified using the NanoDrop 1000 spectrophotometer (Thermo Fisher Scientific, Waltham, MA) and electrophoresis in the 0.8% agarose gel.

The DNA samples of domesticated barley were genotyped using PCR markers distinguishing loss-of-function alleles of the brittleness genes *Btr1* and *Btr2*. The markers were amplified using allele-specific primer pairs Btr1f 5’-CCGCAATGGAAGGCGATG-3’ / Btr1r 5’-CTATGAAACCGGAGAGGC-3’ (∼200 bp fragment, presence - *Btr1* / absence - *btr1*) and Btr2f 5’-AATACGACTCACTATAGGGTTCGTCGAGCTCGCTATC-3’ / Btr2r 5’-GTGGAGTTGCCACCTGTG-3’ (∼ 160 bp fragment, 15 bp deletion in the *btr2* allele). PCR reactions (1 x PCR buffer, 0.1 M primers, 1 U Taq polymerase, 100 ng DNA) were incubated in the PTC DNA Engine thermocycler (Bio-Rad, Hercules, CA, USA) under the following conditions: 95°C for 3 min; 30 cycles of 95°C for 20 s, 60°C for 30 s, 72°C for 1 min; 72°C for 5 min.

### Design of the enrichment assay and SNP calling

To resequence barley genotypes, we designed a custom target enrichment assay, which included 666 loci implicated in the candidate domestication pathways in barley and other species and 1000 neutral loci covering all barley chromosomes to attenuate effects of the biased selection (**Supplementary Fig. 1; Supplementary table 2**). Among the selected loci were known barley genes implicated in regulation of flowering time, development of meristem and inflorescences, tillering, seed dormancy, and carbohydrate metabolism; barley homologs of flowering genes from the other grass species, such as Brachypodium and rice (Higgins *et al.*, 2010); and barley homologs of 259 Arabidopsis genes characterized by the development-related gene ontology (GO) terms that have been confirmed experimentally (**Supplementary table 3**). The barley homologs were extracted from the following sources: the NCBI UniGene dataset (ftp://ftp.ncbi.nih.gov/repository/UniGene/Hordeum_vulgare), IBGSC High and Low confidence genes (IBGSC, 2012), and the HarvEST unigene assembly 35 (http://harvest.ucr.edu). See **Supplementary Methods** for the detailed description of the gene selection and design of the capture baits.

The SNP calling pipeline consisted of three modules: quality control and filtering of Illumina read libraries; mapping the reads to the custom reference; and extracting and filtering both variant (SNP) and invariant sites (**Supplementary Fig. 2**), implemented in a series of bash scripts using standard bioinformatics tools (**Supplementary Methods**).

To determine the ancestral status, the SNPs were genotyped *in silico* in two *Hordeum* species, *H. bulbosum* and *H. pubiflorum* using the Hordeum exome Illumina datasets and the aforementioned bioinformatics pipeline (Mascher *et al.*, 2013a). Alleles that were identical in both species were assigned as ancestral.

*De facto* captured regions were defined as those with the depth of coverage ≥ 8 in at least one of the samples, which was determined using bedtools v.2.16.2, vcftools v.0.1.11 and R (Danecek *et al.*, 2011; Quinlan, 2014). Functional effects of the SNPs were predicted using SnpEff 3.6b using the custom CDS coordinates (Cingolani *et al.*, 2012). The coordinates were determined on the target genomic contigs based on the Spidey predictions (Wheelan *et al.*, 2001).

### Population structure

The population structure was explored using principal component analysis (PCA), Bayesian clustering algorithms (STRUCTURE and INSTRUCT) and a maximum-likelihood (ML) phylogenetic analysis. The subset of putatively neutral SNPs based on the SnpEff flags with minor allele frequency > 0.05 and missing data frequency < 0.5 was selected. The vcf files were converted into the ped format using tabix utility of Samtools and PLINK 1.9 (Chang *et al.*, 2015). The SNPs in high LD (r^2^ > 0.99) were pruned using PLINK. The PCA was performed using smartpca utility of the EIGENSOFT software 5.0.2 (Patterson *et al.*, 2006).

The fastSTRUCTURE software (Raj *et al.*, 2014) was applied with 20 iterations for a predefined number of populations (K). The optimal K for wild barley was chosen to represent the model with maximum marginal likelihood tested for K from 2 to 25. The output matrices were summarized using CLUMPAK (Kopelman *et al.*, 2015), reordered and plotted using an in-house R script. The INSTRUCT tools, which extends the STRUCTURE model to include selfing (Gao *et al.*, 2007), due to very high computational intensity, was ran on 10 randomly drawn subsamples of 1000 SNP markers for five independent chains.

The geographic centers of the populations were calculated as a median of the latitude and longitude of the genotypes comprising the populations. The vector geographic map dataset was downloaded from Natural Earth repository and manipulated in R (http://www.naturalearthdata.com).

The ML phylogeny rooted to *H. bulbosum* and *H. pubiflorum* outgroup species was estimated from the genome-wide SNP dataset using GTRCAT model with Lewis’ ascertainment bias correction to account for the absence of invariant sites in the alignment and the majority-rule tree-based criteria for bootstopping (autoMRE_IGN) implemented in RAxML 8.2.8 (Stamatakis, 2014). The highly admixed wild genotypes were excluded from the input dataset since gene flow between the genotypes may lead to inaccurate placement of the admixed accession on the bifurcating phylogenetic tree. The trees were manipulated using Dendroscope 3.5.7 (Huson and Scornavacca 2012).

### Identification of domestication sweeps

The putative signatures of selection related to domestication were identified using several complementary tests – the diversity reduction index (π_wild_ / π_domesticated_, DRI), Fay&Wu’s H_norm_ (Zeng *et al.*, 2006) and the composite likelihood ratio (CLR) test. The π and H_norm_ statistics were calculated for the individual loci and sliding 10 cM windows (step 1 cM) using mstatspop software, which account for missing genotypes in the data, with 1000 permutations (release 0.1b 20150803; http://bioinformatics.cragenomica.es/numgenomics/people/sebas/software/software.html). A sum of segregating and invariant sites was used to normalize the π values. The software SweeD 3.3.2 (Pavlidis *et al.*, 2013) was used to calculate the CLR test of Kim and Stephan (2002) as expanded by Nielsen *et al.,* (2005), who implemented the use of the genome-wide allele-frequencies as a reference SFS as opposed to the SFS derived from the neutral standard model. The CLR tests were calculated separately for wild and domesticated subsets from the unfolded SFS of individual loci containing at least four SNPs at two grid points across each locus. The genome-wide reference SFS was calculated using SweeD’s “-osfs” flag and provided for the CLR calculations for the individual loci.

To estimate statistical thresholds of the CLR neutral distribution, following Nielsen *et al.* (2005), we simulated 1000 datasets assuming a standard neutral model without recombination using the coalescent simulation software ms with the number of segregating sites (S) and the number of samples (n) as the input parameters describing the wild and domesticated barley populations (Hudson, 2002). Variation of the CLR and in the simulated neutral datasets was assessed using the SweeD with the the threshold to reject neutrality at 99^th^ percentile of the neutral CLR values. The variation of H_norm_. in the same neutral datasets were assessed using msstats software (https://github.com/molpopgen/msstats) and the threshold was chosen as 99,9^th^ percentile to minimize the number of false positives due to the likely deviation of barley demographic history from the standard model. For the DRI, the top 95^th^ percentile was used as a cutoff value following Liu *et al.* (2017). Significance of the overlaps between the tests were estimated using hypergeometric test in R.

## Ancestry of domesticated barley genomes

To estimate ancestry of the domesticated barley loci, we calculated pairwise ML distances between each wild and domesticated genotypes separately for each locus (i.e. individual contig in the mapping reference; in total 39,6 million comparisons) using the GTRGAMMA model in RAxML 8.2.8 (Stamatakis, 2014). If an allele in a domesticated genotype had a smallest ML distance with a population-specific wild allele, this wild allele was deemed ancestral for this locus in this domesticated genotype. The cases where a domesticated barley allele was equally distant to wild barley alleles found in several populations represent the instances of incomplete lineage sorting and therefore were not accounted in a cumulative ancestry of a genotype. A sum of all loci with assigned origin in a single domesticated accession sorted by the locus name we termed an ancestry palette. The ancestry palettes of the individual accessions were pair-wise correlated using Jaccard index-based similarity measure (J) implemented in R:

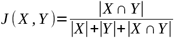

where X and Y are vectors of individual elements of the ancestry palettes, i.e. concatenated locus name and corresponding ancestral population (e.g. locus1_population1) in a pair of genotypes. The ancestry similarity of 1 means identical ancestry palettes and of 0 means that no loci originate from the same population in a pair of accessions. Ancestry similarity heat-maps were visualized using ‘heatmap.2’ function of the ‘gplots’ R package. The scripts and the accompanying files used for the analyses are available in an online repository at https://github.com/artempankin/korffgroup.

## Results

### >500,000 SNPs discovered by the targeted re-sequencing assay

A total of 433 barley accessions, including 344 wild and 89 domesticated barley genotypes were analyzed in this study. To maximize diversity, the barley genotypes were selected to cover the entire range of wild barley habitats in the Fertile Crescent and to represent the whole variety of domesticated barley lifeforms from various breeding programs (**Supplementary Table 1**). Additionally, domesticated barley is classified into the *btr1* (*btr1Btr2*) and *btr2* (*Btr1btr2*) types based on the allelic status of the spike brittleness genes *Btr1* and *2;* independent mutations in either of these genes convert the wild-type brittle spikes into the non-brittle spikes of the domesticated forms (Pourkheirandish *et al.*, 2015). To further verify representativeness of the selected genotypes, we screened for the *Btr* mutations using allele-specific markers. In our genotype set, the *btr1* and *btr2* types were represented by 71% and 29% of the domesticated accessions, respectively (**Supplementary Table 1**).

Illumina enrichment re-sequencing of 23,408 contigs in 433 barley genotypes yielded ∼ 8 billion reads (0.56 Tb of data; deposited at NCBI SRA BioProject PRJNA329198; **Supplementary Note 1** and **Supplementary Table 4**). Cumulatively, the captured regions comprised approximately 13.8 Mbp (**Supplementary Table 5**) and 1.33 Mbp of which resided in the coding regions (CDS). Per sample analysis of the coverage revealed that approximately 87% of the captured regions were covered above the SNP calling threshold and that the between-sample variation was relatively low with the median depth of coverage varying from 45 to 130 (**Supplementary Fig. 3**). The SNP calling pipeline identified 544,318 high-quality SNPs including approximately 190,000 of singletons (**Supplementary Table 5**). Of all the SNPs, 37,870 resided in CDS and approximately 43% of them were non-neutral based on the SnpEff predictions. The CDS were more conserved than the non-coding regions with the average SNP density of 29 and 41 SNPs per Kbp, respectively. 45% of the SNPs could be located on the barley genetic map, whereas for 37% of the SNPs, only the chromosome could be assigned (**Supplementary Fig. 4a**).

### Admixture between wild and domesticated barley

In domestication studies, where patterns of genetic variation are contrasted between wild and domesticated genotypes, it is critical to distinguish these subgroups and exclude genotypes of unclear provenance. The PCA based on the SNP markers revealed two distinct clusters corresponding to the domesticated and wild subspecies with multiple genotypes scattered between these clusters (**Fig. 1a**). fastSTRUCTURE analysis revealed patterns of admixture in 36% and 12% of the domesticated and wild genotypes, respectively, which corresponded to the genotypes intermediate between wild and cultivated barley clusters in the PCA (**Fig. 1b**). Both fastSTRUCTURE and INSTRUCT models produced matching admixture patterns (r^2^ > 0.99) (**Supplementary Fig. 5**). The admixed domesticates did not originate from any specific locality and the admixed wild barley were spread all over the Fertile Crescent indicating that the admixture was not restricted to any particular geographical area (**Supplementary table 1**). These admixed genotypes of ambiguous provenance were removed from the further analyses.

**Figure 1.**
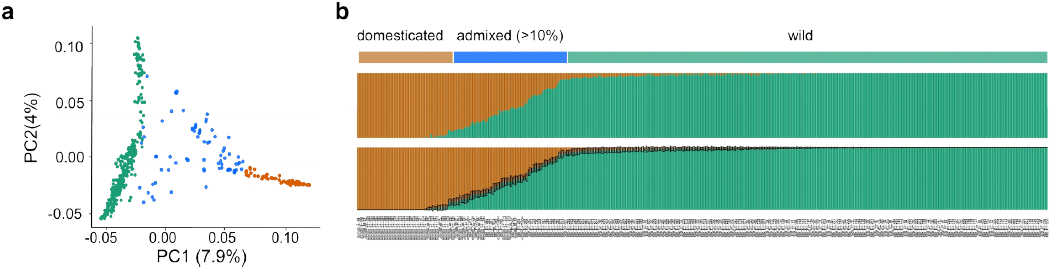
Genome-wide analysis of nucleotide diversity. **(a)** Principal component (PC) analysis of 433 barley genotypes. The first two PC discern subgroups of wild (green), domesticated (orange) barley and admixed (blue) genotypes. A percentage of the total variation explained by the PCs is shown in parentheses. **(b)** Global genetic ancestry of the wild and domesticated barley genotypes as determined by the population structure analysis using fastSTRUCTURE (∼315,000 SNPs) and INSTRUCT (10 random samples of 1000 SNPs) models - upper and lower panels, respectively. Proportions of wild and domesticated ancestral clusters are shown as green and orange vertical bars, respectively. The standard deviations on the INSTRUCT plots are shown as whiskers.

### Footprints of domestication-related selection

Natural selection acting on a beneficial mutation affects various aspects of genetic variation such as allele frequency, nucleotide diversity and linkage disequilibrium at the neighboring regions in a process called selective sweep. To scan for signatures of selective sweeps, which occurred during domestication (hereafter domestication sweep), we performed genome scans using several statistics - the composite likelihood ratio test (CLR), Fay & Wu’s H_norm_, and the diversity reduction index (DRI; π_wild_/π_domesticated_). These statistics explore different patterns of molecular variation and therefore presumably reveal signatures left by selection under different scenarios (Innan and Kim 2004). Both the CLR and H_norm_ statistics are site frequency-based and describe variation of the site frequency spectrum (SFS) at the tested loci. Strong deviations of these statistics from the expected genome-wide values, as tested by coalescent simulations under the neutral scenario, indicate selection. The DRI statistics reveals a severe depletion of nucleotide diversity in the domesticated genotypes at certain loci detected as statistical outliers. The DRI statistics has frequently been used in the selection scans to reveal domestication sweeps in other crop species (Civáň *et al.*, 2015; Huang *et al.*, 2012; Lin *et al.*, 2014; Zhong *et al.*, 2017).

Altogether the scans identified 137 outlier contigs carrying signatures of a selective sweep and 91 of them could be located on the map and covered all barley chromosomes (**Supplementary Table 6**). Of those, 20 contigs (16 mapped locations) were outliers in at least two of the scans (**Fig. 2a**). The overlap between the CLR and H_norm_ scans was relatively high 38% (p-value < 1.0e-07) and the overlaps between the DRI scan and the other two tests were significant but less prominent (8 - 10%; p-value < 0.05) consistent with the previously reported values (Liu *et al.*, 2017).

**Figure 2.**
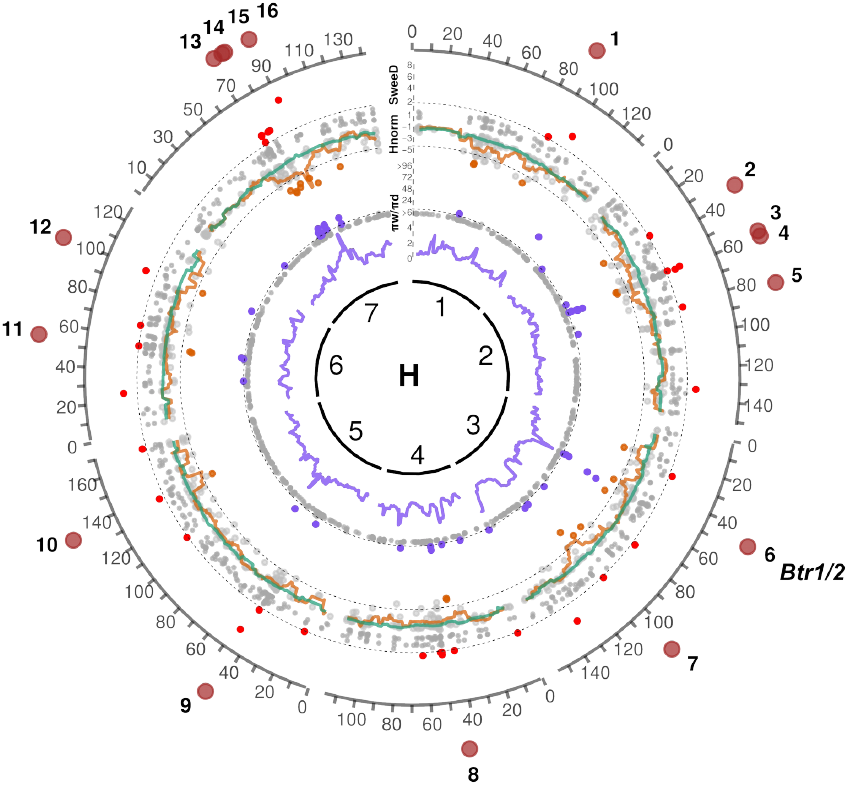
Genomic signatures of domestication selective sweeps and examples of parallelism in the candidate targets of selection in various crops. Genome scans for signatures of selection associated with domestication. The sliding-window and individual-target values are shown as lines and points, respectively. The innermost circle represents barley linkage groups (H) followed by the diversity reduction index (π_wild_/ π_dom_) (violet); the normalized Fay&Wu’s H_norm_ statistics for the wild (green) and domesticated (orange) groups; and the composite likelihood ration statistics (SweeD CLR) for the domesticated group (red). The outlier thresholds are shown by dashed lines and the non-outlier loci are shown as gray dots for all the tests. 16 candidate selected regions supported by at least two of the statistics are shown as brown circles on the outermost layer. *Btr1*/*2* – brittle rachis domestication genes (Pourkheirandish *et al.*, 2015).

Among the top outliers including the loci in the overlaps between the tests were homologs of genes implicated in modulation of the light signaling, circadian clock, and carbohydrate metabolism pathways (**Supplementary Table 6**). None of the candidate domestication genes identified in this study have been functionally characterized in barley, however, putative function can often be inferred from homology.

In our scans, the barley homolog of the light signaling and circadian clock gene *EMPFINDLICHER IM DUNKELROTEN LICHT 1* (*EID1*) had the strongest CLR signal among all the genes (seq375; CLR = 7.13; **Fig. 2c**). In tomato, *EID1* has been implicated in domestication (Müller *et al.*, 2016). The homologs of the Arabidopsis *CULLIN4* (*CUL4*; seq442, AK371672; CUL4, H_norm_=-5.1, CLR = 2.7) and *SUPRESSOR OF PHYA 2* (*SPA2*; seq108, MLOC_52815; DRI = 53), encoding two members of the COP1-CUL4-SPA protein complex implicated in regulation of the light signaling pathway, are other examples where strong signatures of selection have been found in our scans and in another crop species (**Fig. 2b**) (Zhu *et al.*, 2008). In common bean, the homologs of *CUL4* and *CONSTITUTIVELY PHOTOMORPHOGENIC 1* (*COP1*) - a gene encoding another member of the COP1-CUL4-SPA complex, have been independently targeted by selection in two separate domestication events (Schmutz *et al.*, 2014). Two genes of the starch metabolism pathway – the homologs of the alpha- and beta-amylases (AMY, seq669, DRI = 23,; BAM1, seq345, H_norm_ = -5.32) were strong outliers in the DRI and H_norm_ scans. Previous studies discovered reduced variation at the alpha-amylase locus in barley domesticates compared to wild genotypes and hinted at the functional divergence of the wild and domesticated alpha-amylase alleles (Cu *et al.*, 2013; Kilian *et al.*, 2006).

The location of the only known barley domestication locus *Brt1/2* coincided with the domestication sweep 6 on the chromosome 3 and thus the Brt1/2 genes were likely direct targets of selection within the sweep 6 (**Fig. 2a**). The location of selective sweeps 13 – 16 presumably corresponded to the region of depleted diversity on the chromosome 7 discovered by Russell *et al.* (2016), who speculated that the *NUD* gene, controlling the naked (hulless) grain phenotype (Taketa *et al.*, 2004; Pourkheirandish *et al.*, 2015), might have been a direct target of domestication in this region. In our scans, the *NUD* gene itself did not carry a selection signature and thus apparently was not the target of selection at this locus. Indeed, both hulless and hulled genotypes are ubiquitously present in the domesticated barley genepool and thus naked grain phenotype represents an improvement but not domestication trait (Saisho and Purugganan 2007).

### Population structure and ancestry analyses

Next, we investigated whether the local ancestry patterns in cultivated barley genomes could shed light on the phylogeographic history of barley domestication. To this end, we first explored population structure of wild barley genotypes using Bayesian clustering and phylogenetic analyses and then searched within the wild gene-pool for putative ancestral alleles for each locus of the domesticated genotypes using the maximum likelihood approach (see **Materials and Methods**). Following a widely-held assumption, we assumed that any individual genomic locus in a domesticated barley accession has likely descended from a wild population that carry the phylogenetically closest allele and the habitat of this wild population indicates a place of origin of the cultivated barley allele (Civáň *et al.*, 2015; Pourkheirandish *et al.*, 2015). The cases where a putative ancestral allele was not specific to a single wild population were excluded from the analysis to alleviate adverse effects of the incomplete lineage sorting on the ancestry estimates.

Nine wild barley populations were suggested by fastSTRUCTURE, which corresponded to the clearly defined clusters on the phylogenetic tree (**Fig. 3abc; Supplementary Fig. 6; Supplementary Note 2)**. The domesticated barley genotypes branched off as a monophyletic cluster on the phylogram **(Supplementary Fig. 7**). Six wild populations, Carmel and Galilee (CG); Golan Heights (GH); Hula Valley and Galilee (HG); Judean Desert and Jordan Valley (JJ); Negev Mountains (NM); Sharon, Coastal Plain and Judean Lowlands (SCJ), were concentrated in the South Levant and the other three, Lower Mesopotamia (LM), North Levant (NL) and Upper Mesopotamia (UM), occupied large areas of the Northern and Eastern Fertile Crescent. Habitats of the wild populations were distinct with very few immigrants and genotypes of mixed ancestry occurring mostly in the borders of overlapping areas (**Supplementary Figs. 8, 9**). Only 23 wild accessions had a highly admixed ancestry and could not be attributed to any of the nine populations (**Fig. 2c**).

**Figure 3.**
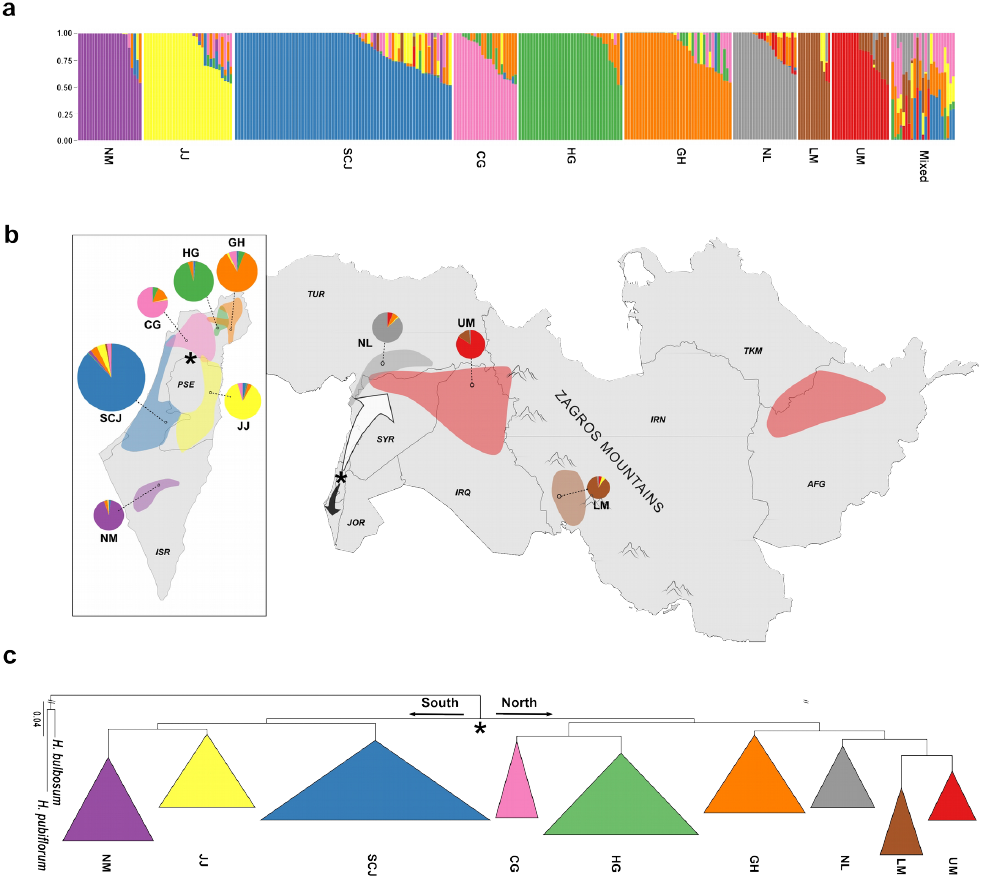
Geographic distribution, structure and phylogeny of nine wild barley populations. The colors correspond to the nine wild barley (*H. vulgare* ssp. *spontaneum*) populations. Carmel & Galilee (CG; pink); Golan Heights (GH; orange); Hula Valley & Galilee (HG; green); Judean Desert & Jordan Valley (JJ; yellow); Lower Mesopotamia (LM; brown); Negev Mountains (NM; magenta); North Levant (NL; gray); Sharon, Coastal Plain & Judean Lowlands (SCJ; blue); Upper Mesopotamia (UM; red). **(a)** Population structure of wild barley as determined by fastSTRUCTURE for K=9. Vertical bars correspond to individual genotypes and colors indicate their membership in the nine subpopulations. **(b)** Distribution of the wild barley populations within the Fertile Crescent. The pie charts represent the ancestral composition of the populations as determined by fastSTRUCTURE and are connected to the geographic centers of population distributions by dashed lines. The size of the pie charts reflects the number of genotypes in the populations. An approximate location of the ancestral wild barley population is shown by an asterisk, and the northward and southward migration routes are indicated by the white and black arrows, respectively. The country codes (ISO 3166) are shown in italics. **(c)** The Maximum Likelihood (ML) phylogeny of wild barley accessions. The clusters were collapsed based on the population assignment. The most ancient population split is indicated by an asterisk. H*. bulbosum* and *H. pubiflorum* were used as distant outgroup species and the length of the outgroup branch was artificially shortened.

Rooting the phylogeny to the outgroup species *H. bulbosum* and *H. pubiflorum* enabled tracing the population differentiation in time. The most ancestral wild population split was located in the north of modern Israel indicating the place where the oldest *H. spontaneum* populations apparently established. Accordingly, wild barley further spread along two routes, the short southern route until the Negev Desert and the longer route to the eastern part of the Fertile Crescent (**Fig. 3ab**). The hierarchy of the populations splits on the phylogram as a function of genetic distance between the clades followed geographical patterns of differentiation and migration of the wild populations away from northern Israel thus indicating the isolation-by-distance pattern in the population structure of wild barley subspecies.

A putative wild ancestor could be assigned for 1,232 loci separately in each domesticated genotype (**Fig. 4a**). 60% of the loci were monophyletic, i.e. a locus that descended from the same wild barley population in all domesticated genotypes, whereas ancestry of 40% of the loci could be traced back to several wild populations across the domesticated genotypes. For further analyses, we separated the dataset into two parts: the candidate domestication loci, to identify geographical origin of the domestication sweep events even without knowing direct targets of selection; and the rest of the genome, which we tentatively termed neutral, to search for the genome-wide signatures of independent founder lineages.

**Figure 4.**
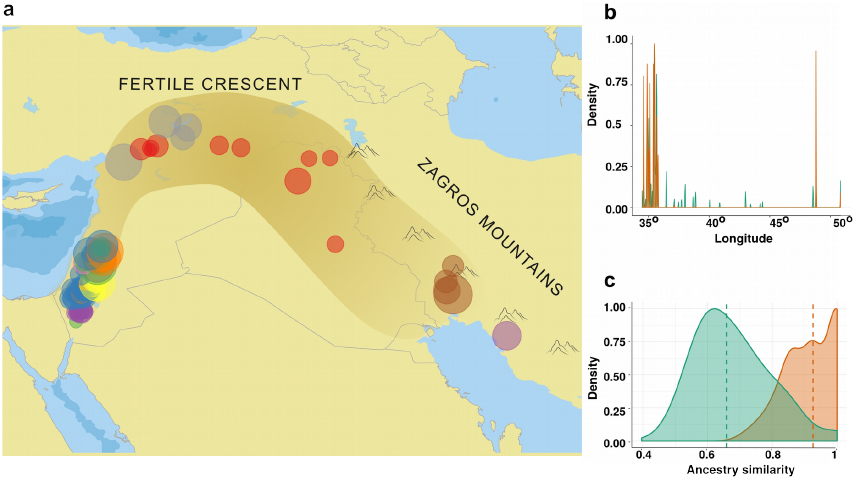
Origin and ancestral compositionof the domesticated barley genomes. **(a)** Geographic distribution of wild barley accessions, which carry ancestral haplotypes of the domesticated barley loci, within the Fertile Crescent. Colors correspond to nine wild barley population as in Fig. 3. **(b)** Longitudinal distribution of the ancestral haplotypes for the neutral (green) and domestication sweep loci (orange). **(c)** Density diagrams of the pairwise ancestry similarity coefficients determined for the neutral (green) and domestication sweep loci (orange).

The distribution of ancestral haplotypes of the neutral loci suggested that wild barley populations from every sampled part of the Fertile Crescent contributed to the domesticated genomes, whereas, in the case of the domestication sweep loci, the longitudinal distribution of the ancestral haplotypes was significantly different (Kolmogorov-Smirnov test p-value < 1e-12) (**Fig. 4ab**). The domestication sweep loci descended from two discernible clusters of wild barley genotypes in the eastern and western parts of the Fertile Crescent (**Fig. 4b**). Intriguingly, the proportions of individual contributions of the ancestral wild populations did not noticeably differ between the domesticated genotypes hinting at a single highly admixed lineage at the root of domesticated barley (**Supplementary Fig. 10**).

To gather further evidence on this hypothesis, we quantified the similarity of the ancestry patterns in the genomes of the domesticates by pair-wise correlating the sorted ancestry palettes of individual accessions. The ancestry palettes of the neutral loci were only moderately similar (median 0.64), which means that, for multiple loci, the patterns of ancestry were not consistent across the domesticated genotypes (**Fig. 4c**). Apparently this dissimilarity of the ancestral patterns at multiple loci resulted from a gene flow, which randomly shuffled alleles descending from different ancestral wild populations between the domesticated genotypes. By contrast, the ancestry palettes of the domestication sweep loci were remarkably similar (median 0.96) across the domesticated genotypes compared with the neutral genome, indicating that the randomizing effect of gene flow was considerably weaker at the genomic regions maintaining the domestication syndrome (**Fig. 4c**). It is noteworthy that this difference between the similarity of the ancestry palettes in the neutral and domestication sweep loci did not arise from the unbalanced number of loci in the subgroups (**Supplementary Fig. 11**). In both subsets, the heat maps of the ancestry similarity did not reveal any clear-cut patterns, e.g. presence of several discernible clusters, which could be interpreted as a signal of distinct founder lineages (**Supplementary Fig. 12**). We therefore propose a following demographic scenario of barley domestication – independent lineages of barley cultivation, which originated from several founder events in the east and the west of the Fertile Crescent, have been combined by means of gene flow into a single admixed (proto)domesticated lineage, which was at the root of the domesticated barley genotypes (**Supplementary Fig. 13**). Continuing gene flow between wild and domesticated subspecies further randomized the ancestry patterns of the modern domesticated genotypes particularly at the selectively neutral loci.

## Discussion

### Selection scans reveal multiple domestication sweeps in barley

Our understanding of the genes and traits that constitute the barley domestication syndrome is extremely limited. Here, the selection scans identified a domestication sweep at the *Btr1/2* locus, which modulates spike brittleness - the only studied example of a crucial domestication trait (*sensu* Abbo *et al.*, 2014). Besides, we discovered multiple novel candidate domestication genes, that are implicated in regulation of light signaling circadian clock, and carbohydrate metabolism pathways. It is noteworthy that the domestication loci detected in this study may be only a representative sample of the truly selected loci. Some loci might have experienced selection regimes leaving signatures that escape detection, confounded with the effects of demography or that are missed because of the gaps in certain regions of the genetic and physical maps (Teshima *et al.*, 2006; Mascher *et al.*, 2013b, 2017).

Intriguingly, we found examples of genes carrying selection signatures both in barley and other crops suggesting convergence of domestication-related selection on homologous developmental pathways and protein complexes in different crop species. The most prominent examples were a circadian clock gene *EID1*, which is a domestication gene in tomato (Müller *et al.*, 2016), and genes SPA and CUL4 encoding components of the E3 ubiquitin-ligase COP1-CUL4-SPA, which was targeted by domestication of common bean. The COP1-CUL4-SPA complex is a critical part of the far-red light signaling, photoperiod and circadian clock pathways (Zhu *et al.*, 2015).

This finding adds to the growing evidence that components of the circadian clock, light signaling and shade avoidance pathways were targets of domestication and further adaptation to new environments in various crop species (Faure *et al.*, 2012; Müller *et al.*, 2016; Shor and Green 2016; Zakhrabekova *et al.*, 2012). However, the evolutionary role such modifications play in the domestication syndrome has not been understood. Müller *et al.*, (2016) suggested that modification of the circadian clock was a human-mediated adaptation of cultivated tomato, which was domesticated in the equatorial regions, to long photoperiods of the northern latitudes. In our study, many of the domesticated barley genotypes originated from the same latitude where barley domestication ensued, which makes, in the case of barley, the scenario of adaptation to a latitudinal cline less plausible. An alternative hypothesis might be linked to the fact that many crop plants are cultivated in dense stands that result in dramatic changes in the light environment and, as a consequence, alters plant architecture compared with their wild ancestor species. Therefore, we propose that such common patterns in the crop adaptation to agricultural practices might be the key to understanding the involvement of the modulators of light signaling, circadian clock and shade avoidance pathways in domestication.

### Ancestry of the candidate domestication loci suggests multiple domestications

Identification of the candidate domestication genes enables predicting the phylogeorgaphic origin of the domestication sweep events, which together with the surveys using neutral markers revealing the closest wild ancestor of the domesticated populations at the genome-wide level represent two complementary approaches to untangle domestication histories (Badr *et al.*, 2000; Civáň *et al.*, 2015; Huang *et al.*, 2012; Matsuoka *et al.*, 2002; Morrell and Clegg, 2007; Poets *et al.*, 2015; Pourkheirandish *et al.*, 2015). Here, the heterogeneous ancestry of the candidate domestication loci provided compelling evidence that multiple domestication sweeps occurred in the background of various founder populations of wild barley. The ancestral populations of the domestication sweep loci were confined to the eastern and western parts of the Fertile Crescent.

The dominant narrative of the barley domestication history has since long revolved around the idea of the two independent domestication lineages originating in the Levantine (west) and Zagros (east) horns of the Fertile Crescent (hereafter east-west model). It stems from the finding suggesting existence of the Occidental and Oriental types of domesticated barley corresponding to the b*tr1* and b*tr2* types, respectively (Takahashi, 1955). Later, the east-west model was strongly supported by the molecular analyses of barley population structure and the geographical distribution of the *btr1* and *btr2* mutations as well as by the archaeological studies (Azhaguvel and Komatsuda, 2007; Morrell and Clegg, 2007; Fang *et al.*, 2014; Morrell *et al.*, 2014; Pourkheirandish *et al.*, 2015; Riehl *et al.*, 2012, 2013; Tanno and Willcox, 2012).

Our findings based on the origin of the domestication sweep loci expand the east-west model by suggesting that not only two but multiple domesticated lineages existed in the past in the Levantine and Zagros clusters of the origin of agriculture. The presence of a third mutation conferring the nonbrittle rachis phenotype of domesticated barley supports this hypothesis (Civáň and Brown, 2017).

Recently, based on the genome-wide SNP genotyping data, Poets *et al.* (2015) presented the mosaic model of barley domestication, which suggests that the genomes of modern barley landraces consist of a mixture of ancestral blocks originating in the five wild barley populations from different parts of the Fertile Crescent (Allaby, 2015; Poets *et al.*, 2015). We found that, in contrast to the domestication sweep regions, the neutral partition of the domesticated barley genomes comprised ancestral blocks that descended from all nine wild barley populations corroborating the mosaic model. Moreover, the ancestral patterns at the neutral loci were not very similar across the genotypes. This indicates that post-domestication the gene flow between the wild and domesticated subspecies and the domesticates themselves continued reshuffling the ancestral blocks in the modern domesticated genotypes, thus erasing the genome-wide signatures of independent domestication lineages. A simulation study demonstrated that, in the case of neutral markers, the gene flow between the independent domestication lineages indeed hinders identification of the founder events (Allaby *et al.*, 2008). By contrast, the domestication sweep loci had nearly uniform ancestry patterns across the genotypes. This exhibits importance of retaining specific ancestral alleles of the domestication genes, which are critical for maintaining the domestication syndrome traits.

Involvement of gene flow in domestication has been documented in other crop species (Huang *et al.*, 2012; Civáň *et al.*, 2013; Hufford *et al.*, 2013). The model of rice domestication is arguably the most vivid example. In rice, two different possibly extinct wild *Oryza rufipogon* populations were ancestors of the *indica* and *japonica* domesticated subspecies, however, the domestication sweep loci originated once in *japonica* and were later introgressed into the *indica* lineage (Fuller, 2011; Huang *et al.*, 2012; Huang and Han, 2015; Choi *et al.*, 2017; but see Civáň *et al.*, 2015). We suggest that, in contrast to the rice model, in the barley domestication history, gene flow was not an isolated event but likely a continuous process, which ensued in the early domestication era and was apparently facilitated by modern breeding. Indeed, the genome of a 6000-year old barley landrace carried signatures of the wild barley introgressions thus confirming instances of the gene flow in the early domesticates (Mascher *et al.*, 2016).

We have yet to understand the exact nature and sequence of demographic events that formed the complex mosaic ancestry patterns during the apparently protracted process of barley domestication. Involvement of several wild populations and abundant continuous gene flow in the process of barley domestication greatly complicates explicit modeling of realistic demographic history of its domestication. What is clear, however, is that it was not constrained to a single center of domestication and involved intensive exchange of the early domesticates between the Neolithic farming communities. Recent evidence predicting migration between the early agriculturalist settlements of the east and the west of the Fertile Crescent hints at a likelihood of such scenario (Lazaridis *et al.*, 2016).

To further unravel barley domestication history characterization of direct targets of selection with the domestication sweep regions is of utmost importance. Our catalog of the candidate domestication loci will facilitate future efforts to characterize novel domestication genes, which modulate yet unstudied aspects of the barley domestication syndrome.

## Author Contributions

A.P. and M.K. conceived and designed the experiments. J.A. and D.B. conducted the enrichment sequencing experiments. A.P. analyzed the data. A.P. and M.K. wrote the manuscript.

## Acknowledgements

We cordially thank Kerstin Luxa, Teresa Bisdorf, Caren Dawidson, Elisabeth Kirst and Andrea Lossow for excellent technical assistance; Eyal Fridman, Hakan Özkan and Benjamin Kilian for barley seeds; and Angela Hancock for critical suggestions on the manuscript. This work was supported by the Max Planck Society and by the Deutsche Forschungsgemeinschaft grants (DFG SPP1530 "Flowering time control: from natural variation to crop improvement") and the Excellence Cluster (EXC1028). A.P. was supported by an IMPRS fellowship from the Max Planck Society.

